# Somatic chromosome pairing has a determinant impact on 3D chromatin organization

**DOI:** 10.1101/2023.03.29.534693

**Authors:** Marta Puerto, Mamta Shukla, Paula Bujosa, Juan Perez-Roldan, Srividya Tamirisa, Carme Solé, Eulàlia de Nadal, Francesc Posas, Fernando Azorin, M. Jordan Rowley

## Abstract

In the nucleus, chromatin is intricately structured into multiple layers of 3D organization important for genome activity. How distinct layers influence each other is not well understood. In particular, the contribution of chromosome pairing to 3D chromatin organization has been largely neglected. Here, we address this question in *Drosophila,* an organism that shows robust chromosome pairing in interphasic somatic cells. The extent of chromosome pairing depends on the balance between pairing and anti-pairing factors, with the anti-pairing activity of the CAP-H2 condensin II subunit being the best documented. Here, we identify the zinc-finger protein Z4 as a strong anti-pairer that interacts with and mediates the chromatin binding of CAP-H2. We also report that hyperosmotic cellular stress induces fast and reversible chromosome unpairing that depends on Z4/CAP-H2. And, most important, by combining Z4 depletion and osmostress, we show that chromosome pairing reinforces intrachromosomal 3D interactions. On the one hand, pairing facilitates RNAPII occupancy that correlates with enhanced intragenic gene-loop interactions. In addition, acting at a distance, pairing reinforces chromatin-loop interactions mediated by Polycomb (Pc). In contrast, chromosome pairing does not affect which genomic intervals segregate to active (A) and inactive (B) compartments, with only minimal effects on the strength of A-A compartmental interactions. Altogether, our results unveil the intimate interplay between inter-chromosomal and intra-chromosomal 3D interactions, unraveling the interwoven relationship between different layers of chromatin organization and the essential contribution of chromosome pairing.

## INTRODUCTION

Eukaryotic chromatin is arranged within a 3D nuclear context, the architecture of which is intimately associated with transcriptional regulation. Several hallmarks of chromatin organization have been described, from chromosome territories and compartments to topologically associating domains (TADs) and chromatin loops (for a recent review see^1^). Chromosome pairing represents an additional level of genome organization. It is well established that, after DNA replication, sister chromatids remain tightly paired along their length (reviewed in^2–4^). In addition, homologous chromosomes extensively pair in germ cells during meiosis and, to a lesser extent, in somatic cells (reviewed in ^5–7^). Prominent somatic pairing of homologues was first reported in Diptera^8^. In particular, it is well established that *Drosophila* homologous chromosomes form extensive end-to-end pairing interactions throughout interphase^7, 9, 10^. These interactions are important for gene regulation, and pairing is found in somatic cells throughout most stages of development^7, 11–20^. The extent of somatic homologue pairing outside of *Drosophila* is still a matter of debate. However, in response to specific cellular processes and/or developmental cues, localized homologue interactions have been identified in multiple species, including mammals^5–7^. Notably, in humans, extensive chromosome pairing appears to be associated with disease^6, 7^.

Somatic chromosome pairing in *Drosophila* occurs at “buttons” located frequently across the genome, resulting in tight association of homologues and sister chromatids^11–14, 17^. Pairing buttons overlap genomic *loci* consisting of numerous architectural protein binding sites (APBSs), and it is thought that combinations of architectural proteins cooperate to modulate pairing^11, 15, 19^. In this regard, an RNAi screen identified multiple pairing promoting and anti-pairing genes^21^, many of which are well evolutionarily conserved^22^. Among these is CAP-H2, a subunit of the condensin II complex that binds to APBSs ^7, 9–11, 23–25^ and has a well characterize function as an anti-pairer^7, 11, 26^. Here we identify Z4, also known as putzig, as a strong anti-pairer that interacts with CAP-H2 and is required for its binding to chromatin. Z4 has been implicated in several functions including Notch signaling, the ecdysone response, germ cell development, and nucleosome remodeling^27–29^.

How inter-chromosomal pairing interactions impact the other features of 3D chromatin organization is not well understood. Here, we have addressed this question in *Drosophila*, which, besides pervasive homologue pairing, shows some other distinct organizational features. On the one hand, *Drosophila* TADs have been shown to best represent close range compartmental interactions, forming compartment domains^30^. In addition, unlike mammals, *Drosophila* chromatin does not arrange itself into CTCF loops^30^. In fact, in *Drosophila*, CTCF is one amongst several other architectural proteins that co-occupy pairing APBSs buttons^11, 12, 15^. On the other hand, *Drosophila* chromatin is organized into additional features; including gene loops and Polycomb (Pc) loops^11, 26, 31–33^. Gene loops are represented by intra-genic interactions in Hi-C, the intensity of which correlates with transcription elongation^11^. Pc loops represent high intensity point-to-point interactions anchored by Polycomb^26, 32, 33^.

To analyze the contribution of chromosome pairing to 3D chromatin organization, we have performed Hi-C experiments under changing pairing conditions. Hi-C data contain information on chromosome pairing, but extracting this information is challenging. In this regard, we recently developed a pipeline for identifying chromosomal pairing from the orientation of ligation between homologous fragments in Hi-C^11^. This approach creates 1D signal tracks that detail the intensity of Hi-C derived pairing (hd-pairing) across the genome, resulting in the identification of paired *loci* at high genomic resolution. To alter pairing interactions, we took advantage of the fast and reversible response to 3D chromatin organization that, like previously described in mammalian cells^34^, *Drosophila* cells undergo upon hyperosmotic cellular stress. In *Drosophila* cells, it was shown that hyperosmotic stress reduces nuclear volume, promotes rapid and reversible chromatin condensation, and induces delocalization of architectural proteins into insulator bodies^35–37^. However, how these changes impact 3D chromatin organization is not known. Our data show that hyperosmotic stress induces full chromosome unpairing and, concomitantly, reduces gene loops and Pc loops, affecting segregation of active (A) and inactive (B) compartments only weakly. These effects, which are rapidly recovered upon stress removal, depend on Z4/CAP-H2 even at regions where Z4/CAP-H2 are not present, suggesting that they are triggered by chromosome unpairing. Along the same lines, depletion of Z4/CAP-H2 increases pairing and reinforces gene loops and Pc loops. Altogether these results show the interwoven relationship between the different layers of 3D chromatin organization and unveil the essential contribution of somatic chromosome pairing.

## RESULTS

### Z4 interacts with CAP-H2 to promote chromosome unpairing

CAP-H2 is one of many architectural proteins present at pairing buttons. Notably, analysis of ChIP-seq data for 16 other known architectural proteins showed exceptionally high co-occupancy of CAP-H2 bound *loci* by Z4 (Fig. 1a, OR=274.7). Indeed, ChIP-seq analysis revealed that essentially all CAP-H2 occupied *loci* are also occupied by Z4 (Fig. 1b, c) and that both proteins preferentially localize at the TSS of transcriptionally active genes (Fig. 1d). Furthermore, co-immunoprecipitation experiments confirm interaction between CAP-H2 and Z4 (Fig. 1e). Interestingly, Z4 is a C2H2 zinc-finger protein with potential DNA binding activity, suggesting that it might be involved in CAP-H2 recruitment to chromatin. To test this possibility, we performed knockdown experiments treating cells with dsRNA against Z4, which decreased Z4 protein levels by approximately 80% (Fig. 1f). In these experiments, we used cells treated with dsRNA against LacZ as control. We then performed CAP-H2 ChIP-seq experiments which revealed that depletion of Z4 results in strongly decreased CAP-H2 occupancy (Fig. 1g), concomitant with the knockdown level. These results suggest that Z4 interacts with CAP-H2 and mediates its binding to chromatin. Next, we performed Hi-C experiments to determine chromosome pairing. CAP-H2 has a well characterized anti-pairer function in *Drosophila*^7, 11, 26^. Counterintuitively, it was previously reported that, despite this anti-pairing activity, pairing is high at *loci* bound by CAP-H2 and increases upon CAP-H2 depletion^11^. As a matter of fact, we detected higher hd-pairing signal at APBSs co-occupied by CAP-H2 in comparison to APBSs lacking CAP-H2 (see Fig. 3d, below).

**Fig. 1.**
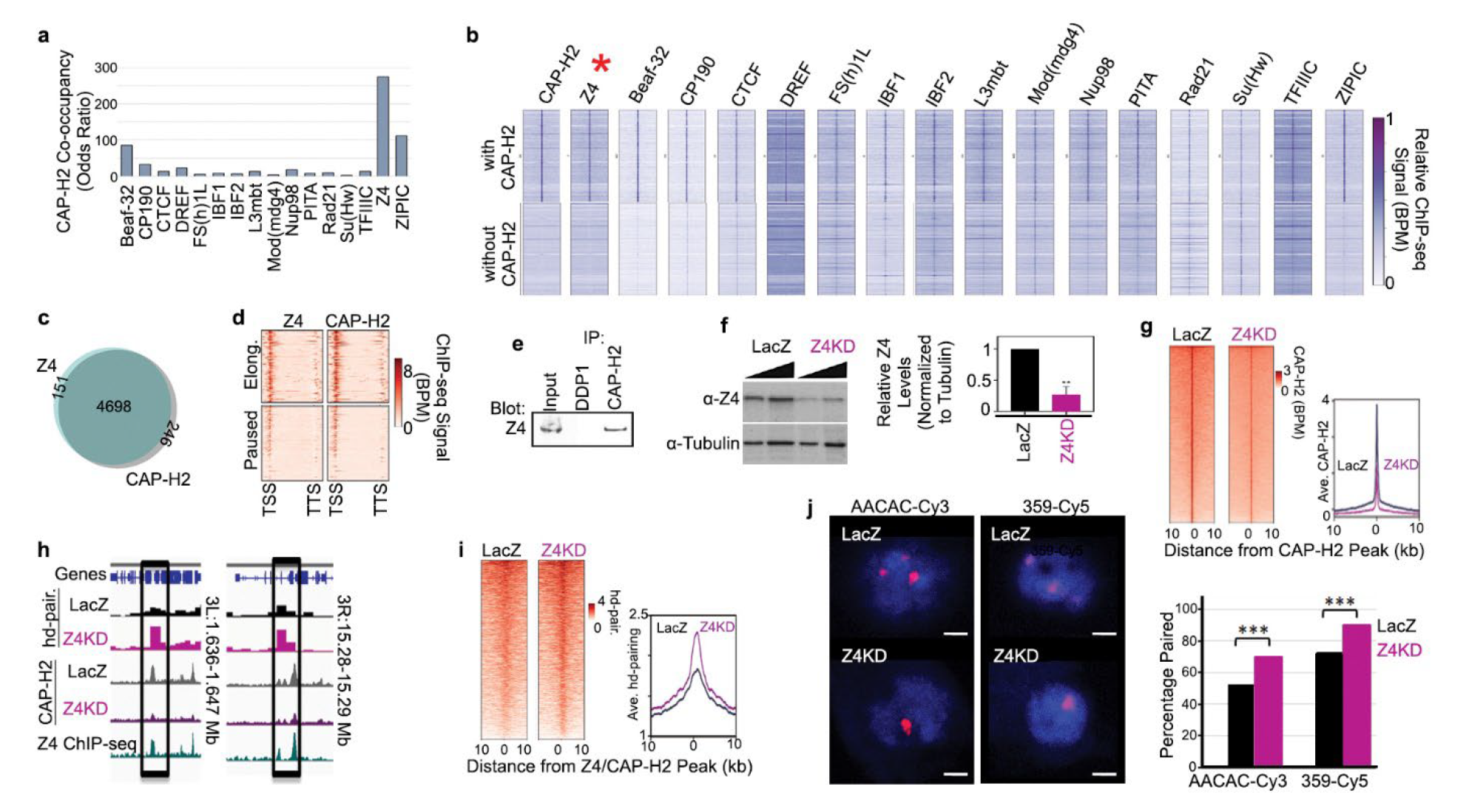
Z4 and CAP-H2 cooperate to control chromosome unpairing. **a)** Odds ratio of the overlap of the indicated architectural proteins ChIP-seq peaks with that of CAP-H2. **b)** Heatmap of ChIP-seq signal for the indicated architectural proteins at APBSs that are occupied by CAP-H2 vs those without CAP-H2. **c)** Venn diagram displaying the number of Z4 and CAP-H2 ChIP-seq peaks that occupy the same *loci*. **d)** Heatmap of ChIP-seq signal for Z4 and CAP-H2 at genes. **e)** Co-immunoprecipitation of Z4 with αCAP-H2 antibodies. Input and IP with αDDP1 antibodies are shown for reference. **f)** Left: WB of Z4 levels in control dsLacZ and Z4 knockdown (Z4KD) cells. Tubulin is shown as loading control. Right: Quantification of the results shown in the right. Z4 protein levels relative to dsLacZ and normalized by the Tubulin control are shown. Results are the average of 3 independent experiments. Error bars are S.D..** indicates p-value < .001; two-tailed Student’s t-test. **g)** Left: Heatmap of CAP-H2 ChIP-seq signal in dsLacZ and Z4KD cells. Right: average profile of the ChIP-seq signal. **h)** Example *loci* showing the hd-pairing signal and CAP-H2 ChIP-seq signal in dsLacZ and Z4KD cells. Z4 ChIP-seq signal in dsLacZ cells is shown to highlight the overlap with CAP- H2. **i)** Left: Heatmap of hd-pairing signal in dsLacZ and Z4KD cells at Z4/CAP-H2 binding sites. Right: average profile of the hd-pairing signal. **j)** Left: FISH analysis in dsLacZ and Z4KD cells for AACAC and 359 (in red), two *loci* commonly studied in conjunction with pairing. DNA was stained with DAPI (in blue). Scales bars correspond to 5μm. Right: Percentage of cells where the *loci* were paired. n = 118 for dsLacZ and n = 124 for Z4KD. *** indicates p-value < .001; two tailed Fischer exact test.

Considering that Z4 strongly co-localizes with CAP-H2 and is required for its binding to chromatin, we anticipated that depletion of Z4 will also increase pairing. Indeed, we observed that, in comparison to control dsLacZ cells, Z4 knockdown cells show increased hd-pairing signal at Z4/CAP-H2 binding sites (Fig. 1h, i). We then chose two *loci*, AACAC and 359, which are commonly used for measuring pairing by Fluorescence *in situ* hybridization (FISH)^21^. We performed FISH for these *loci* and, indeed, confirm increased chromosome pairing in Z4 knockdown cells (Fig. 1j).

Altogether these results suggest that CAP-H2 and Z4 cooperate to inhibit chromosome pairing.

Next, we performed ATAC-seq experiments to test if loss of Z4, and consequently CAP-H2, alters APBSs occupancy. We performed paired-end sequencing to be able to determine Tn5 hypersensitive sites (THSS) as well as fragments consistent with the sizes of nucleosomes^38, 39^ (Fig. 2a). We observed high ATAC-seq signal at APBSs in dsLacZ cells with little to no change after Z4 depletion (Fig. 2b, c). Examining this further, we performed footprint analysis for several architectural proteins and still found little to no change after Z4 depletion (Fig. 2d). These results suggest that increased pairing detected upon Z4 depletion does not correlate with increased APBSs occupancy.

**Fig. 2.**
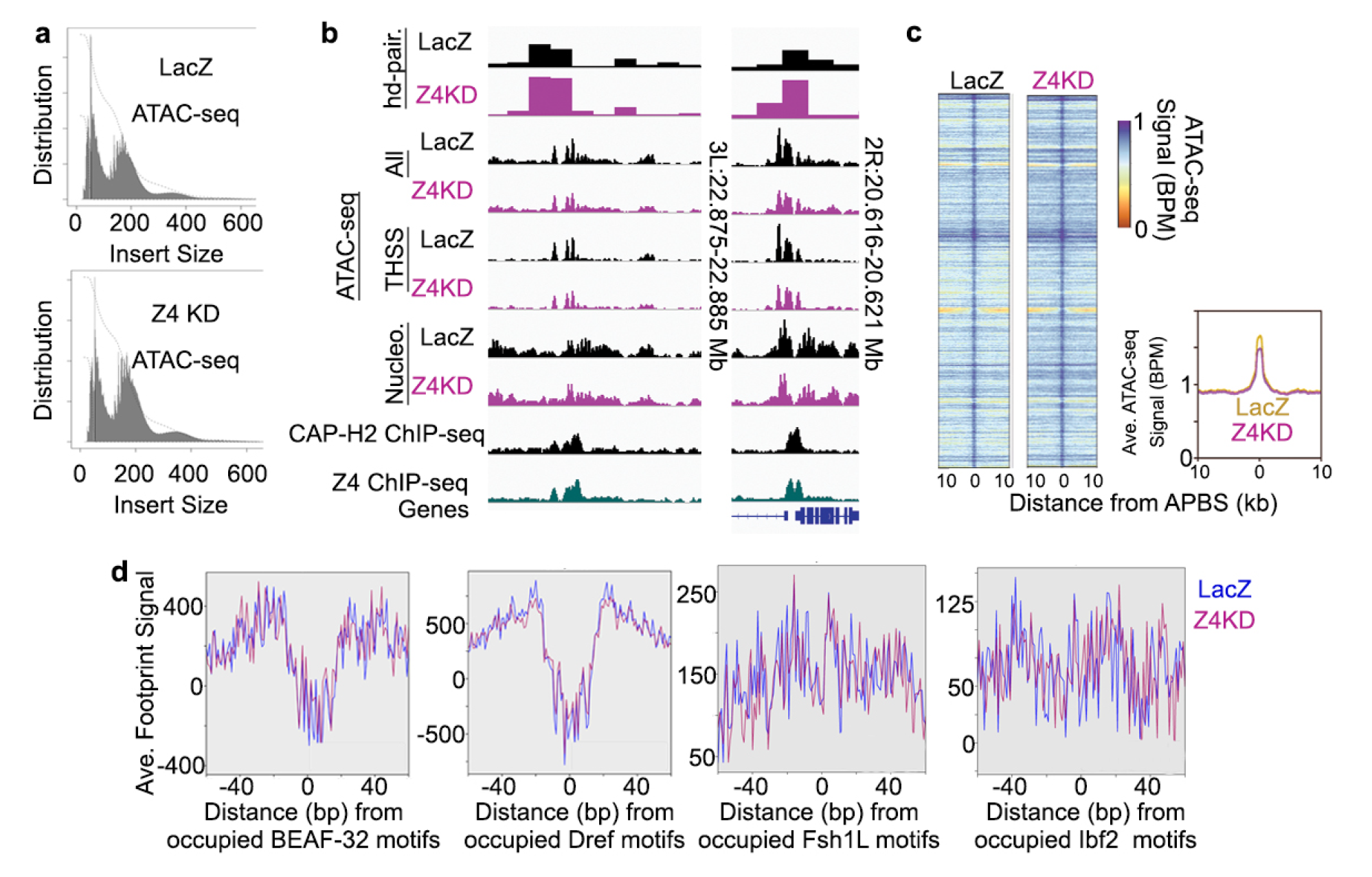
Z4 depletion does not alter occupancy at APBSs. **a)** Distribution of insert sizes obtained from ATAC-seq in dsLacZ and Z4KD cells. **b)** Examples of *loci* with increased hd-pairing signal in Z4KD cells but unchanged ATAC-seq signal. Total signal (All), and reads separated by fragment size into transposase hypersensitive sites (THSSs) and mononucleosomes (Nucleo) are shown to indicate little to no change in each. CAP-H2 and Z4 ChIP-seq signals are also shown. **c)** Heatmap (left) and average profile (right) of ATAC-seq signal at APBSs in dsLacZ and Z4KD cells. **d)** TOBIAS footprints in dsLacZ and Z4KD cells at architectural protein binding sites overlapping Z4/CAP-H2 target *loci*.

### Hyperosmotic stress disrupts chromosome pairing in a Z4/CAP-H2-dependent manner

Chromatin features are highly responsive to environmental stress^34, 40, 41^. However, the impact of stress conditions on chromosome pairing has been largely ignored. To address this question, we performed Hi-C experiments in *Drosophila* cells subjected to NaCl-induced hyperosmotic stress (300 mM NaCl) for 1 hour and after recovery in isotonic conditions for 1 additional hour (Fig. 3a). As previously described^37^, hyperosmotic stress significantly decreased nuclear size (Extended Data Fig. 1a). After hyperosmotic stress, Hi-C contact maps display visible changes in contact intensity (Extended Data Fig. 1b), which correspond to increased long-range (off-diagonal) signal (Extended Data Fig. 1c). The increased interaction range is likely due to the decrease in nuclear volume, thereby constricting the physical space of chromatin. Indeed, osmostress resulted in increased inter-chromosomal (e.g. chr2-chr3) interactions (Extended Data Fig. 1d, e), suggesting that chromosome territories are less distinct. After 1 hour of recovery time, the interaction distance profiles reverted to what was seen in the control (Extended Data Fig. 1b, c) and inter-chromosomal interactions decreased to normal levels (Extended Data Fig. 1d, e), suggesting that broad-scale changes to chromatin architecture in response to cellular hyperosmostress are acute and reversible. A similarly fast and reversible response to osmostress was reported in mammalian cells^34^.

**Fig. 3.**
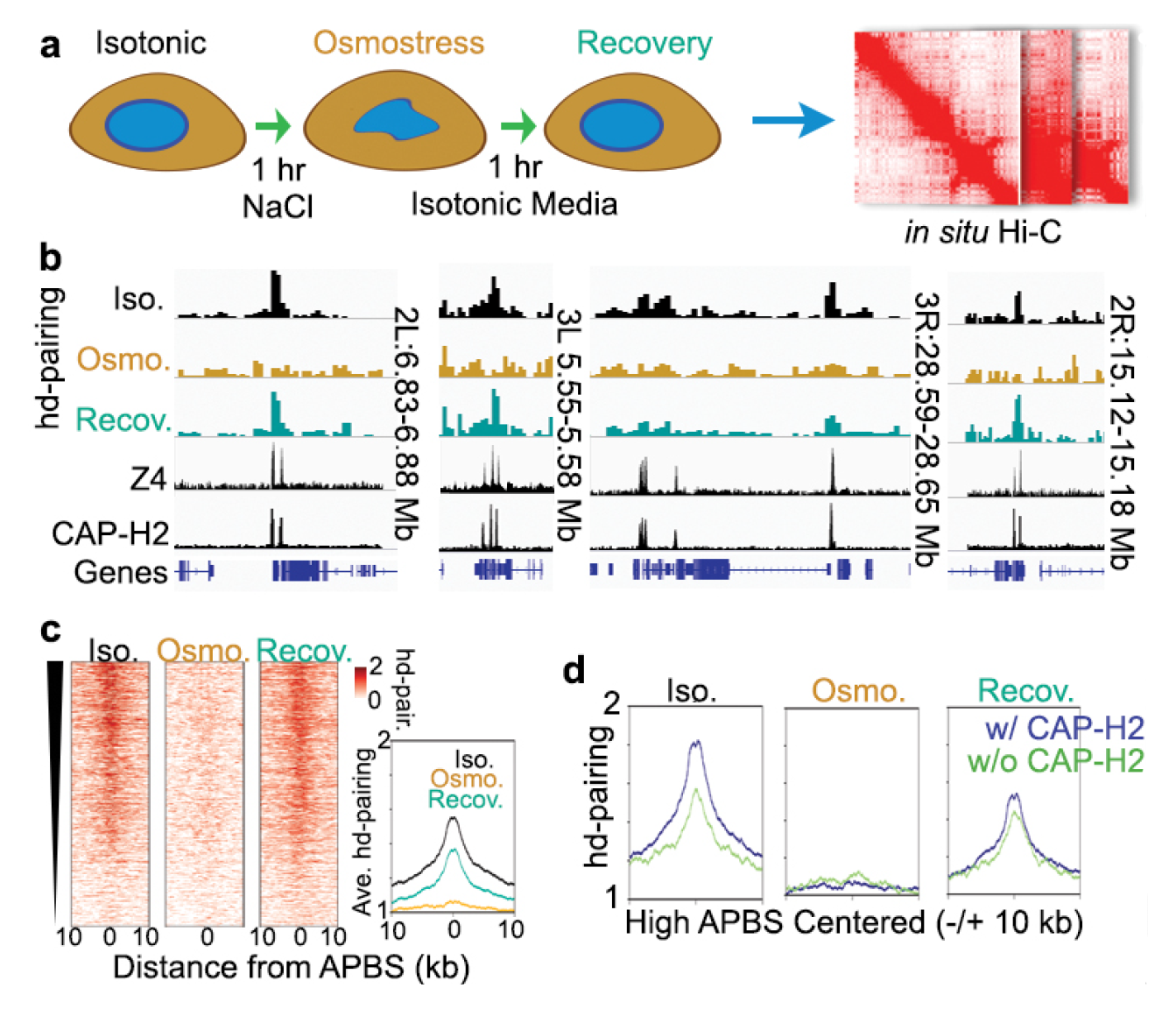
Hyperosmotic stress disrupts chromosome pairing. **a)** Overview of experimental design including hyperosomotic stress conditions along with recovery in isotonic media. **b)** Example *loci* showing the hd-pairing signal under isotonic, osmostress, and recovery conditions. CAP-H2 and Z4 ChIP-seq signals are also shown. **c)** Left: Heatmap of the hd-pairing signal at architectural protein binding sites (APBS) under isotonic, osmostress, and recovery conditions. Right: average profile of the hd-pairing signal under isotonic, osmostress, and recovery conditions. **d)** Average hd-pairing signal at ABPSs that are bound by CAP-H2 (blue) or missing CAP-H2 (green) under isotonic, osmostress, and recovery conditions.

Next, we analyzed the effects of osmostress on chromosome pairing, finding a wide-spread loss of hd-pairing signal at APBSs upon osmostress (Fig. 3b, c). After recovery, pairing was reestablished, but without reaching the original levels (Fig. 3b, c). We then examined separately the effects at APBSs co-occupied by Z4/CAP-H2 in comparison to APBSs lacking Z4/CAP-H2. In isotonic conditions, we detected higher hd-pairing signal at APBSs co-occupied by Z4/CAP-H2 compared to APBSs lacking Z4/CAP-H2 (Fig. 3d, left). Pairing was lost in both categories during hyperosmotic stress (Fig. 3d, middle), but in recovery conditions, pairing was fully restored to APBSs that lack Z4/CAP-H2, but only partially restored to those co-occupied by Z4/CAP-H2 (Fig. 3d, right). This indicates that Z4/CAP-H2 partially inhibits the re-establishment of pairing during recovery.

We also analyzed the effects of Z4 depletion, and thus of CAP-H2, on chromosome unpairing in response to osmostress. As mentioned above, Z4 depletion increases chromosome pairing in isotonic conditions (Fig. 1g, 1h and 4a-c). We observed that, like in control dsLacZ cells, osmostress reduces pairing in Z4 depleted cells (>Fig. 4a-c). However, while osmostress results in nearly complete loss of pairing in control dsLacZ cells (Fig. 4a-c), it was still detected in Z4 knockdown cells to a level that was similar to what is found in control dsLacZ cells under isotonic conditions (>Fig. 4a-c), indicating that loss of pairing during hyperosmotic stress is mediated through Z4/CAP-H2. In addition, we note that, upon recovery from osmostress, pairing is higher than what was originally found in Z4 knockdown cells (Fig. 4a-c). This over-recovery is distinct from what we found in control dsLacZ cells, in which pairing is unable to fully recover (Fig. 3c and 4a-c). This indicates that the presence of Z4/CAP-H2 inhibits pairing during recovery after hyperosmotic stress, and that the absence of Z4/CAP-H2 allows pairing sites to more robustly reassert themselves.

**Fig. 4.**
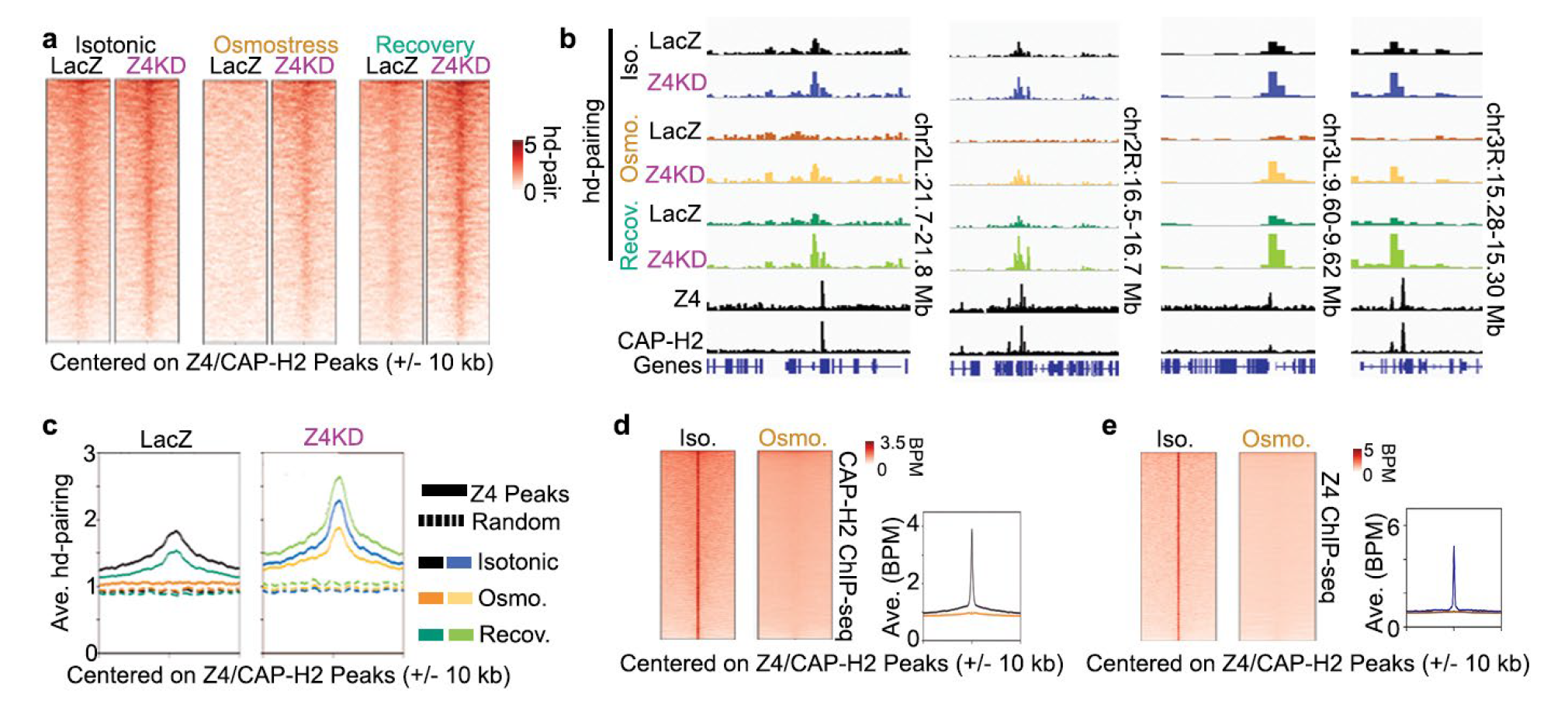
Loss of pairing during osmostress is dependent on Z4/CAP-H2. **a)** Heatmap of hd-pairing signal at Z4/CAP-H2 peaks in dsLacZ and Z4KD cells under isotonic, osmostress, and recovery conditions. **b)** Example *loci* showing hd-pairing signal in dsLacZ and Z4KD cells under isotonic, osmostress, and recovery conditions. Z4 ChIP-seq and CAP-H2 ChIP-seq signal in isotonic conditions are also shown. **c)** Average profiles of hd-pairing signal at Z4/CAP-H2 peaks in dsLacZ and Z4KD cells under isotonic, osmostress, and recovery conditions. Random peaks (dashed lines) are shown to indicate the average background signal at non-Z4/CAP-H2 target *loci*. **d)** Left: Heatmap of CAP-H2 ChIP-seq signal under isotonic and osmostress conditions at CAP-H2 sites found in dsLacZ cells under isotonic conditions. Right: average profile of CAP-H2 ChIP-seq signal. **e)** Left: Heatmap of Z4 ChIP-seq signal under isotonic and osmostress conditions at Z4 sites found in dsLacZ cells under isotonic conditions. Right: average profile of Z4 ChIP-seq signal.

Altogether, these results show that hyperosmotic stress induces fast and reversible chromosome unpairing that is controlled by Z4/CAP-H2 as part of the osmostress response. However, intriguingly, ChIP-seq analyses show a widespread loss of Z4/CAP-H2 peaks upon osmostress (Fig. 4d, e), suggesting that Z4/CAP-H2-mediated unpairing induced by osmostress does not require their stable binding to APBSs.

### Chromosome pairing has a strong impact on gene-loop interactions but affects compartmentalization only weakly

Results reported above show that Z4 depletion and hyperosmotic stress have opposite effects on chromosome pairing, with increases upon Z4 knockdown and decreases in osmostress. Next, we analyzed the effects of these changes on other layers of 3D chromatin organization, namely A/B compartmentalization and gene loops.

Compartments are architectural features that span long distances^1, 30, 42^. We identified compartments at 10 kb resolution by principal component analysis (PCA), the resultant eigenvector denoting A and B compartments that correspond to active and inactive regions, respectively. We found that both Z4 depletion and osmostress result in little to no change to the genomic intervals assigned to A and B compartments (Extended Data Fig. 2a), which are still immediately visible (Fig. 5a). Indeed, if we account for distance-effects by distance-normalization, the resultant maps are visually similar to those of the control conditions (Extended Data Fig. 2b). Separate from compartment position, we also examined compartment strength by saddle analysis, finding only minimal changes (Fig. 5b). We found that the strongest interactions within the A compartment (AA) slightly increase upon Z4 depletion (Fig. 5b, c), while they are weakly diminished in osmostress conditions and revert to initial level in recovery conditions (Fig. 5b, c). On the other hand, the weak B compartment interactions remain mostly unchanged both after Z4 depletion and in osmostress (Fig. 5b, c), except for a weak increase after recovery from osmostress (Fig. 5b, c).

**Fig. 5.**
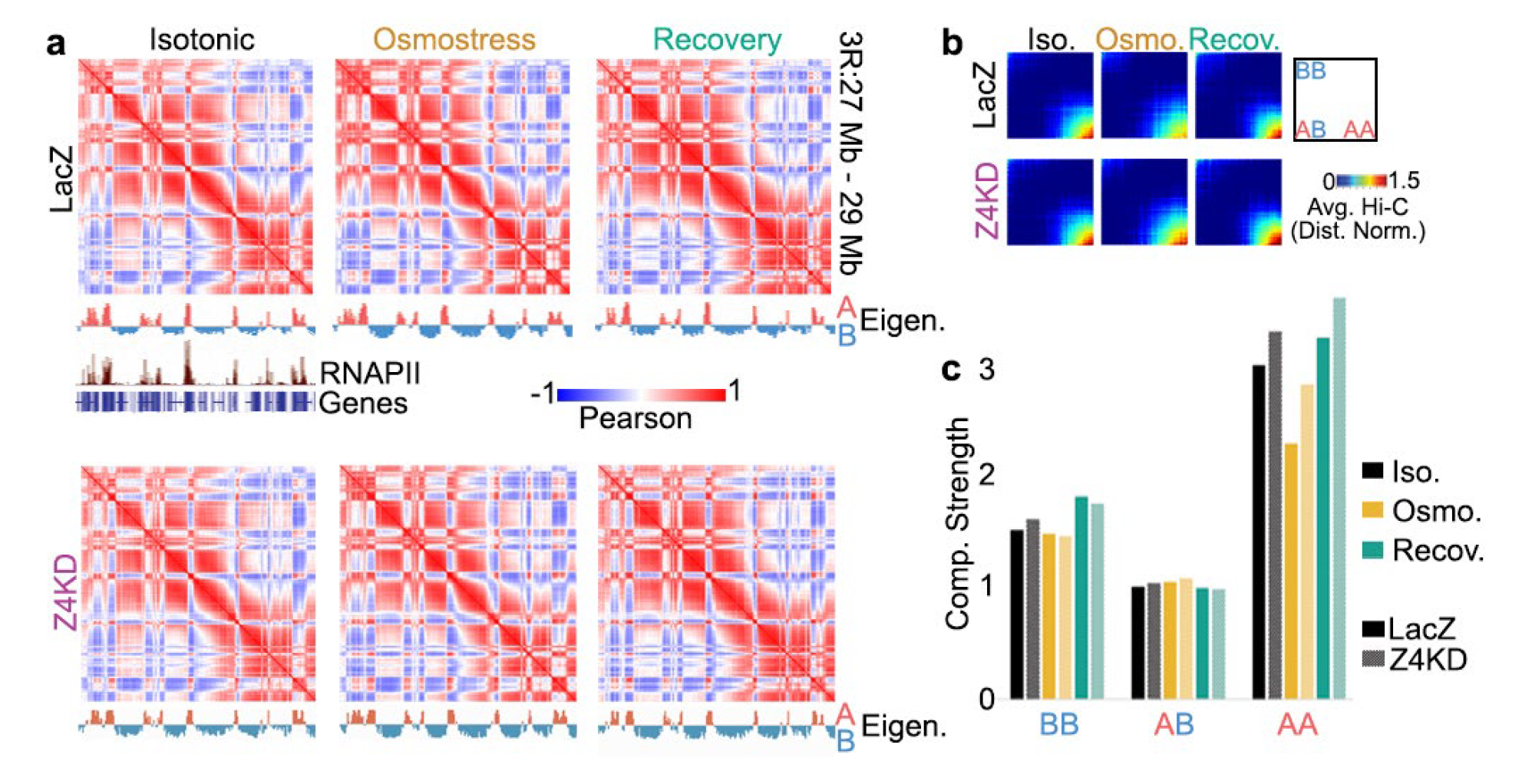
Chromosome pairing affects A/B compartmentalization only weakly. **a)** Pearson correlation matrix of the Hi-C signal in dsLacZ and Z4KD cells under isotonic, osmostress, or recovery conditions. Eigenvector track of the A and B compartments is shown below each. RNAP II ChIP-seq signal in dsLacZ cells under isotonic conditions is also shown. **b)** Saddle plot analysis of compartments in dsLacZ and Z4KD cells under isotonic, osmostress, or recovery conditions. **c)** Compartment strength quantification in dsLacZ and Z4KD cells under isotonic, osmostress, or recovery conditions.

In contrast to compartments, which are largely resilient to changes in chromosome pairing, we observed strong effects on gene-loop interactions. It was previously shown that genes have high intra-genic interactions, and form contacts between the transcription start site (TSS) and the transcription termination site (TTS), otherwise known as gene-loops^11, 30, 43–45^. In *Drosophila*, gene-loops correlate with transcription elongation^11^. We categorized genes by RNAPII ChIP-seq pausing index and, indeed, we detect higher gene-loop interactions on elongating genes compared to paused genes (Fig. 6a, e). We observed increased intra-genic interactions after Z4 depletion, with differences particularly pronounced at elongating genes (Fig. 6a-d). Increased gene-loop interactions occurred specifically at the 5’ and 3’ ends of genes compared to the gene-body (Fig. 6a-d), which matches with the location of Z4/CAP-H2 at the TSS of genes (Fig. 1d). On the other hand, we found that, opposite to Z4 depletion, gene-loops were largely abolished under hyperosmotic stress (Fig. 6a, b), resulting in widespread loss of TSS-TTS interactions (Fig. 6d), especially at elongating genes (Fig. 6e). We also observed that, upon recovery from osmostress, gene-loop interactions were fully restored to their pre-stress level (Fig. 6c-e). Altogether these results suggest a correlation between chromosome pairing and gene-loop interactions.

**Fig. 6.**
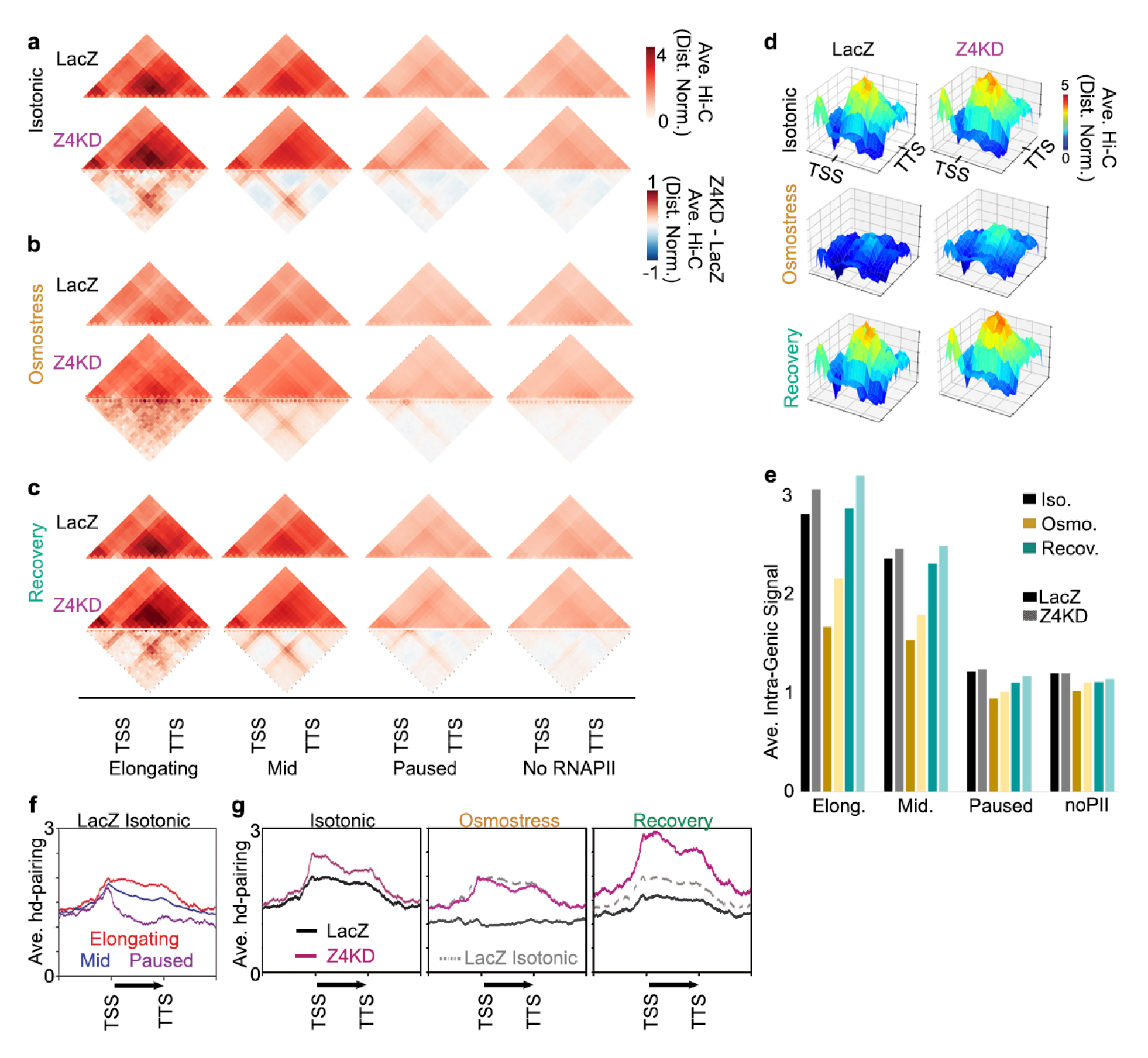
Chromosome pairing impacts intragenic gene-loop interactions. **a-c)** Average distance normalized Hi-C signal in dsLacZ and Z4 KD cells under isotonic **(a)**, osmostress **(b)**, and recovery **(c)** conditions at genes categorized by the RNAPII pausing index. Average difference in intragenic interactions in Z4KD compared to dsLacZ cells for genes categorized by the RNAPII pausing index is also shown. Hi-C signal is distance normalized. **d)** Surface plots with the height indicative of the intensity of the interactions by Hi-C averaged across elongating genes in dsLacZ and Z4 KD cells under isotonic, osmostress, and recovery conditions. **e)** Quantification of the average intra-genic interaction signal in dsLacZ and Z4 KD cells under isotonic, osmostress, and recovery conditions for genes categorized by the RNAPII pausing index. **f)** Average hd-pairing signal in dsLacZ cells under isotonic conditions for genes categorized by the pausing index. **g)** Average hd-pairing signal at elongating genes in dsLacZ and Z4KD cells under isotonic, osmostress, and recovery conditions. Average profile of hd-pairing signal in dsLacZ cells under isotonic conditions (gray dashes) is also shown for comparison.

Next, we addressed the possible causative relation between these two layers of 3D chromatin organization. In this regard, we noticed that pairing interactions occur not only at the TSS, but also throughout the gene-body (Fig. 6f). Indeed, depletion of Z4 leads to increased pairing both at the TSS and throughout the gene body of elongating genes (Fig. 6g, left). The gene-body pairing is lost after hyperosmotic stress in control dsLacZ cells, but is retained in Z4 knockdown cells to levels similar to those observed in control dsLacZ cells under isotonic conditions (Fig. 6g, middle). In recovery conditions, pairing is incompletely reestablished in control dsLacZ cells, but is enhanced throughout the gene in Z4 knockdown cells (Fig. 6g, right). Therefore, despite Z4/CAP- H2 being bound to the TSS (Fig. 1d), they impact pairing interactions not only at the TSS but also across the gene-body. Notably, we observed that, despite partial retention of pairing in Z4 knockdown cells upon osmostress (Fig. 6g, middle), intragenic gene-loop interactions are severely diminished by hyperosmotic stress both in control dsLacZ and Z4 depleted cells (Fig. 6a, b, d, e). These results indicate that, though correlated with, pairing within the gene body is not a downstream consequence of gene-looping.

It was previously reported that gene-looping correlates with RNAPII occupancy^11^. In fact, concomitantly to increased gene-loop interactions, we detected increased polymerase occupancy at TSSs upon Z4 depletion in both elongating and paused genes (Fig. 7a-c). On the other hand, hyperosmotic stress results in widespread decrease of RNAPII ChIP-seq signal (Fig. 7a-c). This effect was most pronounced in elongating genes, in which RNAPII occupancy spreads through the gene-body with only a low level of RNAPII remaining at the TSS (Fig. 7b, c). Weaker effects were detected at paused genes (Fig. 7b, c). These results confirm the correlation of RNAPII occupancy with gene-loops and, hence, with chromosome pairing.

**Fig. 7.**
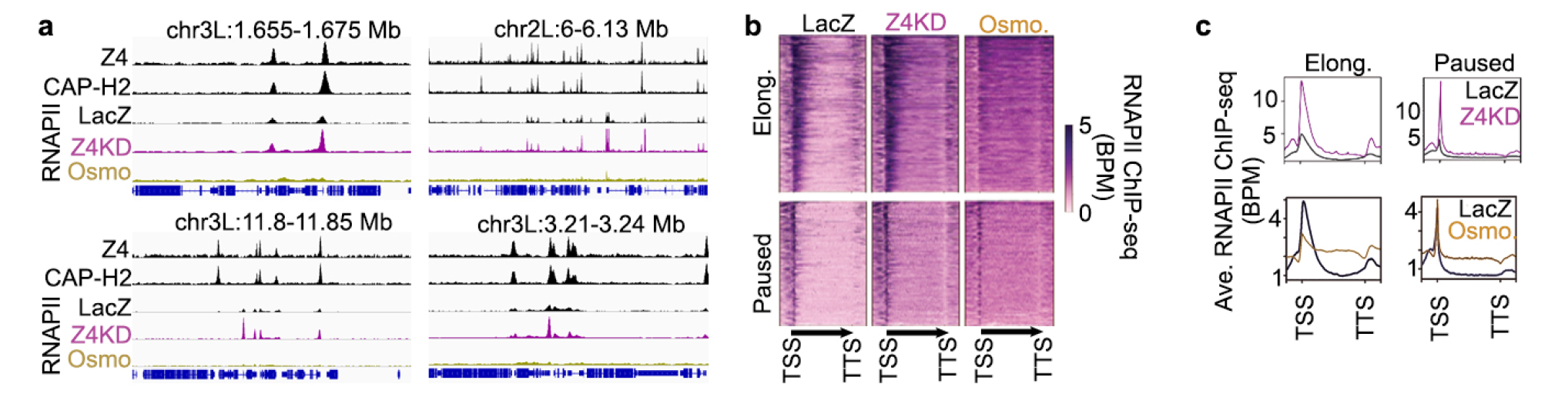
Changes in chromosome pairing alters RNAPII occupancy. **a)** Example *loci* showing the changes in RNAPII ChIP-seq signal in Z4KD cells and under osmostress conditions compared to dsLacZ cells in isotonic conditions. Z4 ChIP-seq and CAP-H2 ChIP-seq signals in dsLacZ cells under isotonic conditions are also shown. **b)** Heatmap of RNAPII ChIP-seq signal in dsLacZ cells under isotonic conditions is compared to those of Z4KD cells under isotonic conditions and dsLacZ cells under osmostress conditions at genes sorted by the pausing index. **c)** Average RNAPII ChIP-seq signal in dsLacZ cells under isotonic conditions is compared to those of Z4KD cells under isotonic conditions and dsLacZ cells under osmostress conditions at genes sorted by the pausing index.

### Chromosome pairing reinforces Pc-loop interactions

Next, we asked if chromosome pairing can impact the interactions at other layers of chromatin organization. To test this, we turned to the few hundred punctate loops anchored by Polycomb (Pc), which are found in *Drosophila* cells^26, 32, 33^ (Fig. 8a, top left). We observed that, concomitantly to reduced pairing, Pc loops signal is lost after osmostress (Fig. 8a, b). Western blot analysis of crosslinked chromatin, indicates that Pc abundance is not affected by osmostress (Extended Data Fig. 3a), suggesting that loss of Pc-looping after osmostress is not associated with reduced levels of chromatin bound Pc. In addition, we observed that, despite the absence of Z4/CAP-H2 binding directly at the loop anchors (Extended Data Fig. 3b), Pc loop signal is increased in Z4 knockdown cells (Fig. 8b, Extended Data Fig. 3c). Moreover, in comparison to control dsLaZ cells, hyperosmotic stress in Z4-depleted cells leads to less pronounced decreases in Pc-looping, with Pc loop signal remaining to nearly the same degree as found in isotonic dsLacZ cells (Fig. 8a, b, c). Examining these regions further, we found hd-pairing buttons nearby corresponding to Z4/CAP- H2 occupancy (Fig. 8a, blue highlight). These hd-pairing buttons are lost upon osmostress and are increased upon Z4 knockdown (Fig. 8a). These results suggest that the strength of Pc-loop interactions is influenced by the extent of pairing at nearby ABPSs *loci*. To further determine the contribution of chromosome pairing to Pc-loop interactions, we examined published haplotype resolvable Hi-C data in a *Drosophila* hybrid cell line^12^. We detect the characteristic punctate Pc loop signal in each of the individual alleles. Intriguingly, this punctate signal is also seen on the inter-allele maps (Fig. 8d, e), indicating that, despite the pairing button being located outside of the Pc loop anchors, part of the Pc loop signal is due to inter-allele interaction^12^. Altogether these results suggest that, even acting at a distance, chromosome pairing impacts other layers of 3D chromatin organization, reinforcing Pc-loop interactions.

**Fig. 8.**
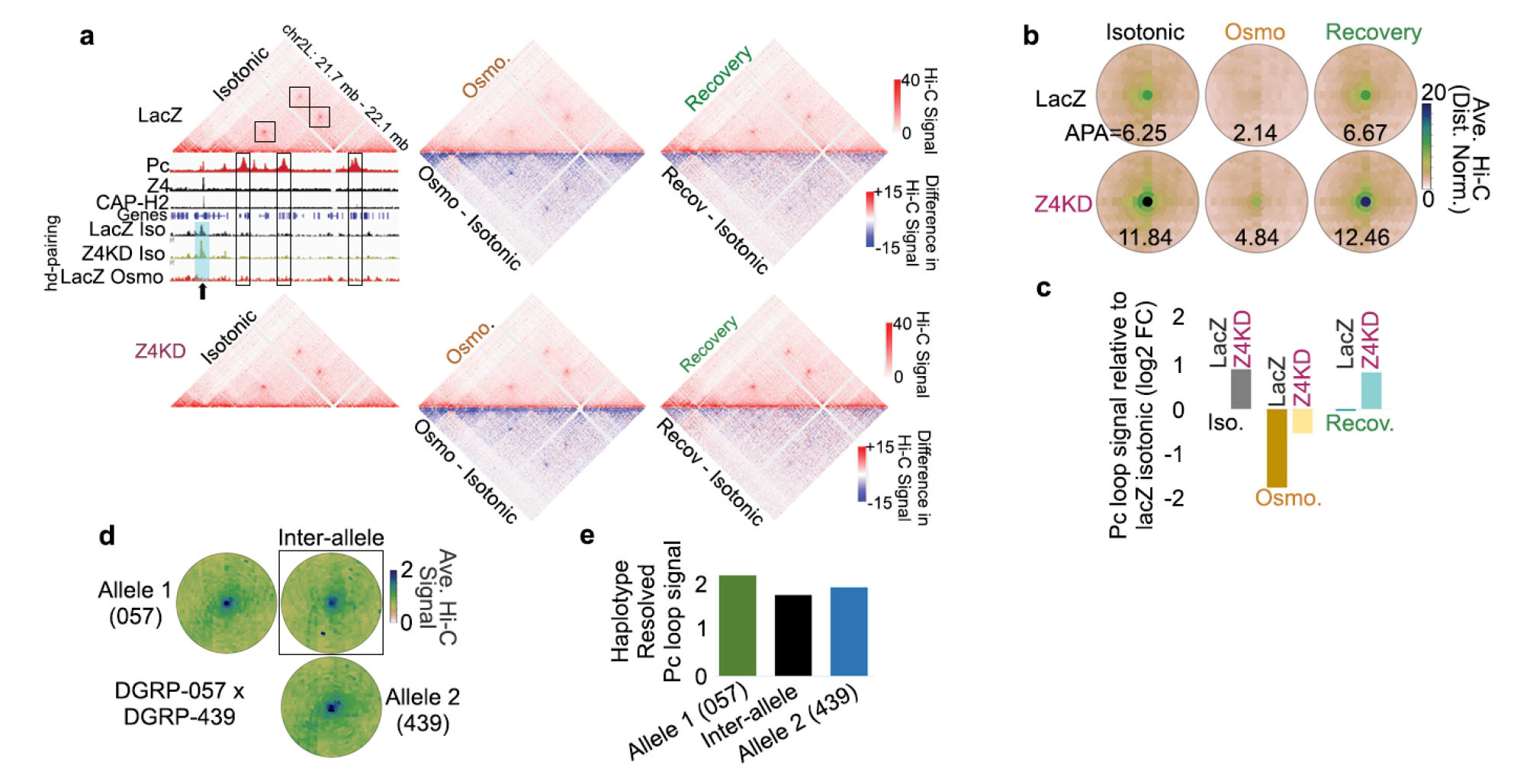
Chromosome pairing reinforces Pc-loop interactions. **a)** Example of Pc loops in dsLacZ and to Z4KD cells under isotonic, osmostress, and recovery conditions. Average difference with respect to isotonic conditions compared to osmostress and recovery conditions in dsLacZ and Z4KD cells are also shown. Pc, Z4 and CAP-H2 ChIP-seq signals in dsLacZ under isotonic conditions are shown to highlight the overlap of Pc but not Z4/CAP-H2 at these loop anchors. hd-pairing signal is shown below to highlight the nearby peak of pairing (arrow) that increases in Z4KD cells and is lost in osmostress conditions. **b)** Average distance normalized Hi-C signal at Pc loops in dsLacZ and Z4KD cells under isotonic, osmostress, and recovery conditions. APA scores for each condition are indicated. **c)** Difference in Pc loop signal for each condition relative to dsLacZ cells under isotonic conditions. Values represent the log2 fold-change of the average distance-normalized Hi-C signal. **d)** Average distance normalized Hi-C signal at Pc loops in the DGRP-057/439 hybrid cell line is shown for the individual 057 and 439 alleles, as well as for the inter-allelic interactions. **e)** Quantification of the average Pc loop on each allele as well as inter-allele haplotype-resolvable Hi-C signal.

## DISCUSSION

Here, we have studied the contribution of somatic chromosome pairing to 3D chromatin organization using *Drosophila* as a model system. It is well known that *Drosophila* chromosomes undergo robust pairing throughout interphase^7^. Tightly paired *loci* correspond to APBSs^11–15, 17^, where a large number of architectural proteins bind. Chromosome pairing results from the balance between opposing pairing and anti-pairing activities^7^. Multiple factors have been reported to influence pairing^21^. However, only few of them correspond to known architectural proteins that might be directly involved in pairing interactions, with CAP-H2’s role in anti-pairing being the most widely studied^7, 9–11, 23–25^. The difficulty in identifying other proteins directly involved in pairing may be related to the proposed cooperative role of architectural proteins^11, 15, 17, 19^. Here, we identify the architectural protein Z4 as an anti-pairer that works in conjunction with CAP-H2. Z4 interacts with CAP-H2 and binds the same *loci*, while depletion of Z4 causes loss of CAP-H2 occupancy and increases pairing. Counterintuitively, despite their anti-pairing activity, Z4/CAP-H2 binding sites show strong pairing and, in fact, pairing at APBSs bound by Z4/CAP-H2 is higher than at sites lacking Z4/CAP-H2, supporting the hypothesis that anti-pairers target pairing sites to unpair and then become unbound^11^. In this regard, we have observed that cellular hyperosmotic stress, which induces chromosome unpairing in a Z4/CAP- H2 dependent manner, results in the delocalization of Z4/CAP-H2 from pairing buttons, suggesting that their unpairing activity does not require stable association with the original site (Fig. 9). Along the same lines, we observed that APBSs pairing buttons have high ATAC-seq accessibility that is not significantly altered upon Z4 depletion, suggesting that pairers and anti-pairers do not compete for binding. Altogether, these observations support a model by which the unpairing function of Z4/CAP-H2 does not involve stable binding to specific buttons, neither is the consequence of competition with pairers for binding (Fig. 9).

**Fig. 9.**
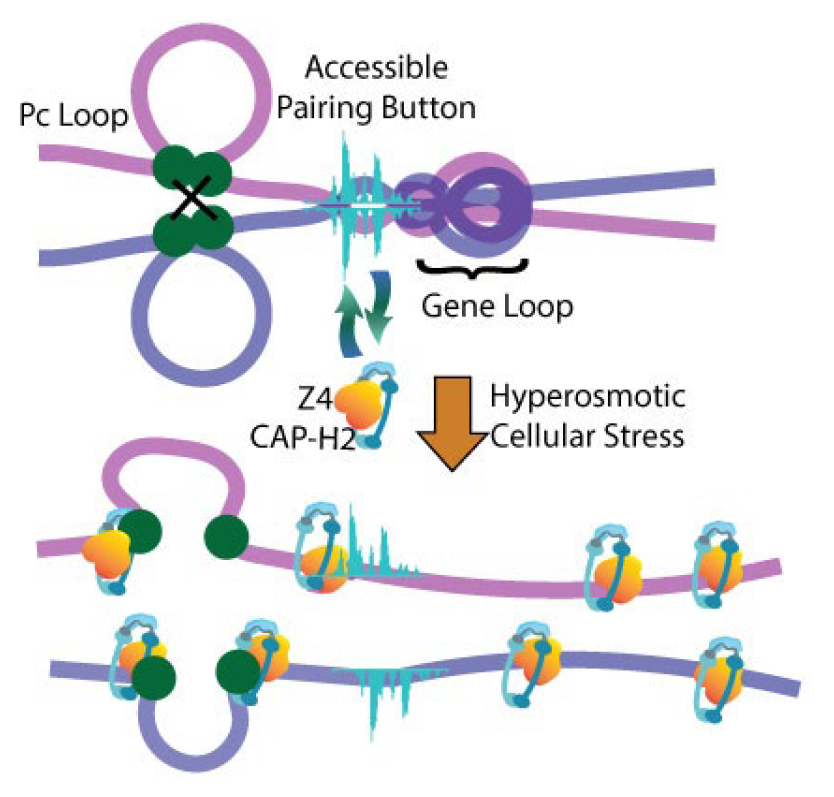
Model for the interwoven relationship of inter-chromosomal pairing interactions with intra-chromosomal layers of 3D chromatin organization. A schematic showing how pairing buttons may enhance nearby organizational features, such as gene-loops and Pc loops. Pairing buttons are ATAC-seq accessible in isotonic conditions and Z4/CAP-H2 targets these *loci*. Under hyperosmotic cellular stress, chromosomes become unpaired in a Z4/CAP-H2-dependent manner. This model would suggest that some Hi-C signal corresponding to gene loops and Pc loops results from the intermingling of paired alleles and that loss of pairing itself weakens these interactions.

Our results indicate that depletion of Z4 and hyperosmotic stress result in opposite changes to pairing and, concomitantly, to gene-loop and Pc-loop interactions, highlighting the interconnectedness between inter-chromosomal and intra-chromosomal 3D interactions (Fig. 9). This is, perhaps, best demonstrated by the altered Pc loop signal observed in Z4 knockdown cells under both isotonic and hyperosmotic conditions, even though Z4/CAP-H2 is not directly bound at these Pc loop anchors. While we cannot rule out other types of effects, a simple explanation is that disruption of normal chromosome pairing impacts other features of chromatin organization. Indeed, inter-allele Pc loop signal provides evidence that pairing is likely contributing to Pc-looping. At the gene level, we observed that pairing has a major impact on intragenic gene-loop interactions. Notably, Z4 depletion increases pairing not only at the TSS region, where Z4/CAP-H2 localize, but all along the gene body, where intragenic interactions accumulate. Furthermore, we note that osmostress in control cells abolishes both intragenic gene loops and pairing, while osmostress in Z4- depleted cells mostly abolishes gene-loops even though pairing is partially retained, suggesting that pairing is not dependent on gene-loop formation.

It has been reported that the extent of intragenic gene-loop interactions correlates with RNAPII occupancy^11, 12, 14^. Consistent with this, we observed that, concomitantly to the loss of pairing and gene loops, the pattern of RNAPII distribution along elongating genes is altered upon hyperosmotic stress, with strongly decreased accumulation at the TSS and increased occupancy along the gene body. Similar results have been reported in mammalian cells^34^, where osmostress induces a global RNAPII run-off from the TSS into the gene body in transcribing genes. Our results also show that Z4 depletion, which reinforces pairing and gene-loop interactions, results in increased RNAPII occupancy at the TSS. Altogether, these results further support the correlation between RNAPII occupancy and gene-loops, extending this correlation to chromosome pairing. Interestingly, chromosomes appear to pair only when they are transcribed and transcription-based models for both somatic and meiotic chromosome pairing have been proposed (reviewed in ^46–49^). In the future, measuring the relative timings of the changes to these features in response to osmostress will help to elucidate which events are upstream of each other.

Like gene loops, segregation of the A-compartment correlates with transcription^11, 12, 14^. However, whether or not the A-compartment and gene loops are maintained by the same mechanism remains unclear. Our results show that, despite the dramatic loss of gene-loops observed at elongating genes upon osmostress, segregation of A and B compartments remains largely unaffected, with only minimal decrease in A-A compartmental interactions. Along the same lines, increased gene-loop interactions observed in Z4 depleted cells coincide with only weak compartmental changes. These results suggest that, at least partially, maintenance of the A-compartment and gene loops relies on distinct mechanisms. The weak effects on compartmentalization observed in *Drosophila* in response to osmostress contrasts with the larger changes reported in mammalian cells^34^, suggesting that compartmentalization in *Drosophila* is more resilient to hyperosmotic stress than in mammalian cells. Whether this is related to the robustness of pairing interactions in *Drosophila* remains to be determined.

Our results also provide insights on the responsiveness of chromatin structure to hyperosmotic stress. It was previously reported that osmostress alters intra-chromosomal 3D interactions in mammalian cells^34^. Here we show that, in addition, cellular osmotic stress in *Drosophila* causes fast and reversible chromosome unpairing (Fig. 9). Osmostress induced chromosome unpairing is likely an active process since it depends on the anti-pairer proteins Z4/CAP-H2. Interestingly, pairing is high at genomic intervals within the active A-compartment (Extended Data Fig. 4a) and Z4/CAP-H2 binding sites are highly enriched in the A-compartment (Extended Data Fig. 4b), suggesting that Z4/CAP-H2 may impact how the A-compartment responds to stress. In fact, we found that hyperosmotic loss of A-A interactions was not as pronounced in Z4 knockdown cells (Fig. 5c). Notably, although there is some debate on the degree of changes to chromatin organization^40, 50^, heatstress also results in a widespread redistribution of architectural proteins as well as shutdown of most transcription^40, 51^. From this point of view, it is conceivable that response to hyperosmotic stress and heatshock may share some common mechanisms^52^. Despite the general loss of transcription, heatshock does lead to induction of several exceptionally short (1-2 kb) Hsp genes^51, 53^, and, notably, we were able to detect increased RNAPII ChIP-seq signal at Hsp70 upon osmostress, with more moderate/varied effects at other Hsp genes (Extended Data Fig. 4c), which is consistent with partially overlapping mechanisms for responding to different types of stress^54^. Altogether, these observations suggest that rapid and reversible changes to chromatin structure is an evolutionarily conserved feature that is observed in response to different types of stress.

In summary, combining osmotic stress and Z4/CAP-H2 depletion, our work demonstrate the interwoven relationship between different layers of 3D chromatin organization, unveiling the essential contribution of chromosome pairing. In this manner, osmotic stress in conjunction with depletion of architectural proteins may serve as an excellent model to derive the interplay between layers of chromatin architecture.

## METHODS

### Antibodies

αCap-H2, αZ4 and αDDP1 are described in ^29, 40, 55^, and ^56^, respectively. The rest of antibodies used were commercially available: αRNAPII (Biolegend, Cat# 664906; RRID:AB_2565554), αTubulin (Millipore, Cat# MAB3408, RRID:AB_94650), αH3 (Cell Signaling, Cat# 9715; RRID:AB_331563), and αPc (Santa Cruz Biotech, Sc-25762).

### Cell culture, RNAi, and salt treatment

*Drosophila* Kc167 cells were cultured in isotonic SFX media. For RNAi knockdown experiments, cells were treated with dsRNAs against Z4 and, as control, against LacZ, using the primers described in Extended Table 1. For Z4 knockdown two different dsRNAs were used in combination: dsRNAVDRC and dsRNAPG obtained using the primers from the Vienna *Drosophila* Resource Center Repository and from Silva-Sousa *et al*.^57^, respectively. Briefly, at day 0 cells were diluted to 10^6^ cells/ml and the dsRNAs added to the media, 20 μg dsRNALacZ/10^6^ cells for the LacZ controls and 10 μg dsRNAPG + 10 μg dsRNAVDRC/10^6^ cells for the Z4 knock-downs. Cells were incubated for 3 days and, at day 3, diluted again to 10^6^ cells/ml and treated with a second dose of dsRNAs as for day 0 for 3 more days. To induce hyperosmotic stress, cells were treated with 300mM NaCl for 1 hour, which is in the range previously used for hyperosmotic stress in *Drosophila* (250mM – 400mM)^35, 36^. Recovery conditions included incubation in the above hypertonic media for 1 hour followed by incubation in isotonic SFX media for 1 hour. Cell viability and p38 phosphorylation status were checked after salt treatment and upon recovery.

### Fluorescence in situ hybridization (FISH)

For FISH, cells were adhered onto lysine coated slides for 1 hour followed by brief washes in PBS for 5 min. Cells were fixed in 4% paraformaldehyde in PBS and washed in 2xSSCT for 5 min at room temperature (RT). Slides were then washed with 2xSSCT/50% formamide for 5 min at RT followed by pre-denaturation in 2xSSCT/50% formamide at 92°C for 2 min and 60°C for 20 min. Hybridization with DNA probes was carried out at 92°C for 3 min by placing DNA probe solution directly onto slides. Slides were then placed in a humidified chamber overnight at RT and mounted with vectashield-DAPI after 2 washes in 2xSSCT at RT. Cells were imaged with Zeiss LSM 780 confocal microscope. Probes used were Cy5-359 (Cy5-GGGATCGTTAGCACTGGTAATTAGCTGC) and AACAC-Cy3 (Cy3-AACACAACACAACACAACACAACACAACACAACAC) from Integrated DNA Technologies (IDT).

### Determination of the effect of hyperosmotic stress on nuclear size

Cells in isotonic media or after hyperosmotic stress were briefly washed with PBS, fixed in 4% paraformaldehyde and stained with DAPI for 15 min. Then, cells were plated on glass slides and subjected to imaging using SPE confocal microscope. Nuclear area measurements were done by using Fiji software.

### Co-IP

Co-IP experiments were performed with total extracts. Briefly, 60 mL of confluent Kc167 were harvested by centrifugation and the pellet was washed three times with PBS. The washed pellet was dissolved in 1 mL of IP Buffer (50mM Tris-HCl pH8, 200mM NaCl, 5mM EDTA, 0.5% NP-40, Protease Inhibitors cocktail), incubated in ice for 30 min, lysed with a Dounce homogenizer B (35- 40 strokes), centrifuged at 3700 g for 15 min at 4°C, and supernatant was aliquoted and stored at −80°C. For IP, 6 μL of rabbit αCAP-H2 and 2 μL of rabbit αDDP1 (as control) were added to the extracts and incubated on a wheel overnight at 4°C. Then, 30 μL of Protein A sepharose beads (GE Healthcare Life Sciences) were added and incubated for 2 hours at 4°C. Beads where recovered by centrifugation and washed 3 times with 1 mL of IP buffer. For elution, beads were resuspended in 2xPLB, β-mercaptoethanol, boiled and spun down, and the supernatants were analyzed by Western Blot (WB).

### ChIP-seq

For Chip-seq, crosslinked chromatin and immunoprecipitation experiments were performed as described previously^58^, except that antibody-bound protein complexes were isolated with protein A sepharose beads (GE Healthcare Life Sciences) washed with RIPA buffer (140mM NaCl, 10mM Tris-HCl pH 8.0, 1mMEDTA, 1% Triton X-100, 0.1% SDS, 0.1% DOC) and shortly blocked with RIPA, 1% BSA. The following antibodies were used for immunoprecipitation: guinea pig αZ4 (2 μL), αrabbit CAP-H2 (6 μL) and mouse αRNAPII (4 μL). For sequencing, single-indexed and dual-indexed DNA libraries were generated using the NEBNextUltra II DNA Library Prep kit from Illumina (New England Biolabs). Single-indexed libraries were sequenced 50 nt single end reads on a HiSeq 2500 (Illumina) getting a minimum of 14 millions reads per sample, and dual-indexed libraries were sequenced 150 nt paired end reads on a NovaSeq 6000 (Illumina) getting a minimum of 25 millions reads per sample. ChIP-seq data was mapped by Bowtie2^59^ to the genome build dm6 with a quality filter ≥10. Mapped reads of replicates were merged together and BPM (bins per million) normalized. Peaks were identified by Macs2^60^. Elongation status was determined by RNAPII ChIP-seq from the pausing index calculated as previously described^11, 61^. Co-occupancy of CAP-H2 with various architectural proteins was calculated by the odds ratio of overlap and non-overlapping peaks from a previously created master list of architectural protein binding sites^26^. Heatmaps and average profiles were generated by deeptools^62^.

### *In situ* Hi-C library preparation

*In situ* Hi-C experiments were performed as previously described^63^ with some modifications. Cross-linked cells were incubated for 30 min in ice in Hi-C lysis buffer (10mM Tris-HCl at pH8, 10mM NaCl, 0.2% NP-40), centrifuged for 5 min at 3500g, resuspended in 190 μL of 1xNEBuffer2 (New England BioLabs [NEB]) and 10 μL of 10% SDS, and were then incubated for 10 min at 65°C. After the addition of Triton X-100 and 15-min incubation at 37°C, nuclei were centrifuged for 5 min at 3500g and were resuspended in 300 μL of 1xNEBuffer2. Nuclei were digested overnight using 400 U of the MboI restriction enzyme (NEB). For fill-in with bio-dATP, nuclei were pelleted and resuspended in fresh 1x repair buffer (1.5 μL of 10mM dCTP, 1.5 μL of 10mM dGTP, 1.5 μL of 10mM dTTP, 37.5 μL of 0.4mM biotin-dATP, 50 U of DNA Polymerase I large [Klenow] fragment in 300 μL 1xNEBuffer2). Ligation was performed for 4 h at 16°C using 10,000 units of T4 DNA ligase (NEB) in 1.2 mL of ligation buffer (120 μL of 10× T4 DNA ligase buffer, 100 μL of 10% Triton X-100, 12 μL of 10 mg/mL BSA, 963 μL of H2O). After reversion of the cross-link, DNA was purified using phenol/chloroform/isoamyl. Purified DNA was sonicated to an average size of 300–400 bp using a Bioruptor (Diagenode; eight cycles of 20 sec on and 60 sec off), and 3 μg of sonicated DNA was used per *in situ* Hi-C library preparation. Briefly, biotinylated DNA was pulled down using 20 μL of Dynabeads MyOne T1 streptavidin beads in binding buffer (5 mMTris-HCl at pH 7.5, 0.5 mM EDTA, 1 M NaCl). End-repair, A-tailing and Illumina adaptors ligation were performed on the beads using NEBNext library preparation end-repair and A-tailing modules (NEB). Libraries were amplified and sequenced in biological replicates, obtaining >200 million useable reads per condition after mapping, deduplication, and merging replicates.

### Hi-C data analysis

Sequenced reads were preprocessed, mapped to the *Drosophila* genome build dm6, filtered for quality score ≥10, and removed of PCR duplicates using HiC-Pro^64^. After initial processing we obtained approximately 200 million useable contacts per condition (Extended Data Table 2), which is enough to resolve structural details of several layers of chromatin organization in the relatively small-sized *Drosophila* genome^30^. Replicates were combined and the resultant maps were visualized with Juicebox^65^. Distance normalization was calculated as the observed value compared to the average value at that distance in the formula (observed+1)/(expected+1). Percent inter-chromosomal interactions was calculated by the number of reads mapping to different chromosomes (e.g. chr2-chr3) v.s. mapping to the same chromosome (e.g. chr2-chr2, chr3-chr3, chr4-chr4, chrX-chrX). Compartments were identified by calculating the eigenvector on the Pearson correlation matrix in 10 kb bins as previously described^30^. Saddle plots were created by sorting interactions by each anchor’s eigenvector percentile (100 quantiles). Compartment strength was calculated by the average distance normalized signal within 5 quantiles in each category. TSS-TTS signal represents distance normalized Hi-C values in the 1 kb binned interaction overlapping the TSS and TTS. Hi-C derived pairing (hd-pairing) profiles across the genome were identified from fragment-level orientation of ligation events in Hi-C as previously described and placed into 1 kb bins^11^. Pc loops were identified by SIP and APA plots were generated by SIPMeta^66^.

### ATAC-seq

ATAC experiments were performed as described^67^. Briefly, nucleus were resuspended in 50 μl Tn5 transposase mixture (2.5 μL Tn5, 25 μL 2XTD buffer, 22.5 μL RNase Free Water), and incubated at 37°C for 30 min. After the reaction, the DNA was purified using the MinElute PCR Purification Kit (QIAGEN). The PCR cycles for the library preparation were determined by qPCR. Following amplification, the size selection of the library was performed using Ampure beads. The dual-indexed libraries were sequenced 50 nt paired end reads on a NovaSeq 6000 (Illumina) getting a minimum of 25 millions reads per sample. ATAC-seq data was trimmed of adapter sequences using cutadapt^68^, aligned to the reference genome dm6 using bowtie2^59^, deduplicated with picard (https://broadinstitute.github.io/picard/), and peaks identified by macs2 with the extsize 147 and nomodel parameters^60^. Tranposase hypersensitive sites (THSSs) and nucleosomes were separately analyzed by categorizing the reads by insert size (<120 bp v.s. 140- 250 bp respectively). Signal was BPM normalized and we tested for differential peaks by MANorm^69^. Footprint analysis was performed using TOBIAS^70^ on Z4 peaks overlapping architectural protein motifs where available in the Jaspar database (i.e. Beaf-32 and Dref) and ChIP-seq summits for the others.

## DATA AVAILABILITY

The accession numbers of the ChIP-seq, Hi-C, and ATAC-seq data generated in this work are GEO: GSE213553; Reviewer Token: adyjesqmxfofleh. APBS ChIP-seq datasets were GSE30740; GSE63518; GSE54529; GSE80702 from^26, 40, 71, 72^. Pc ChIP-seq data set was GSE63518 from ^40^. Haplotype resolved Hi-C for 057×439 hybrid cells was GSE121255^12^.

## ACKNOWLEDGMENTS

We thank Drs. Ting Wu and Jumana AlHaj Abed for helpful discussions during the preparation of this manuscript. We also thank Dr. Victor Corces for antibodies, training and helpful discussions, and Drs. Anja C. Nagel, Asifa Akhtar and Lluisa Espinàs for antibodies. We also acknowledge the help of the IRB Functional Genomics Facility in Hi-C library preparation. This work was supported by the MICIN/AEI 10.13039/501100011033 (BFU2015-65082-P and PGC2018-094538-B-100), ‘FEDER, una manera de hacer Europa’ and the Generalitat de Catalunya (SGR2014-204 and SGR2017-475) (to F.A.); the Spanish Ministry of Economy and Competitiveness (PID2021- 124723NB-C21 and PID2021-124723NB-C22) (to F.P. and E.N.) and the Unidad de Excelencia Maria de Maeztu (MDM-2014-0370) (to UPF). FP and EN are recipients of an ICREA Acadèmia (Catalan Government). S.T. received funding from the European Union’s Horizon 2020 Research and Innovation Programme under the Marie Skłodowska-Curie grant agreement no. 754510. Support was also from the NIH Pathway to Independence Award R00 GM1287671 and the NIH MIRA award R35 GM147467 (to M.J.R.); the content is solely the responsibility of the authors and does not necessarily represent the official views of the NIH.

## EXTENDED DATA

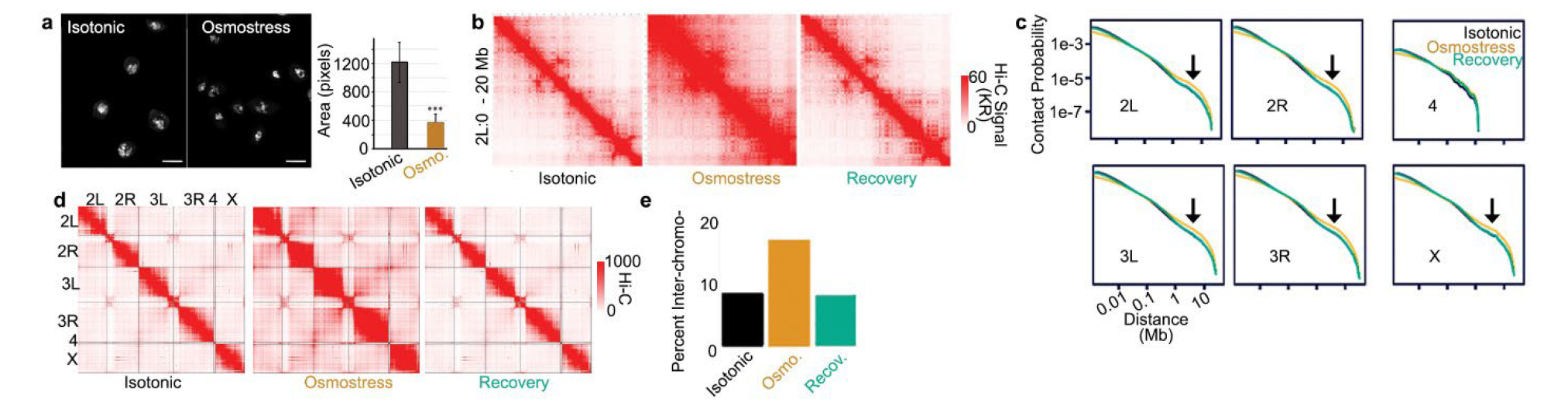
Extended Fig. 1. a) Left: DAPI staining (in white) of nuclei under isotonic and osmostress conditions. Scale bars correspond to 5μm. Right: Nuclear area quantified from DAPI staining. Error bars are S.D..n = 125 for isotonic and n = 128 for osmostressed cells. *** indicates p-value < .001; two-tailed Fisher’s exact test. b) Hi-C maps showing how hyperosmotic stress results in increased off-diagonal signal that returns to normal levels upon recovery from stress. c) Hi-C contact probability along each chromosome under isotonic, osmostress, and recovery conditions. Arrows indicate bump at long range distances in osmostress conditions that is no longer observed under recovery conditions. d) Inter-chromosomal Hi-C maps under isotonic, osmostress, and recovery conditions. e) Percentage of interchromosomal interactions under isotonic, osmostress, and recovery conditions.

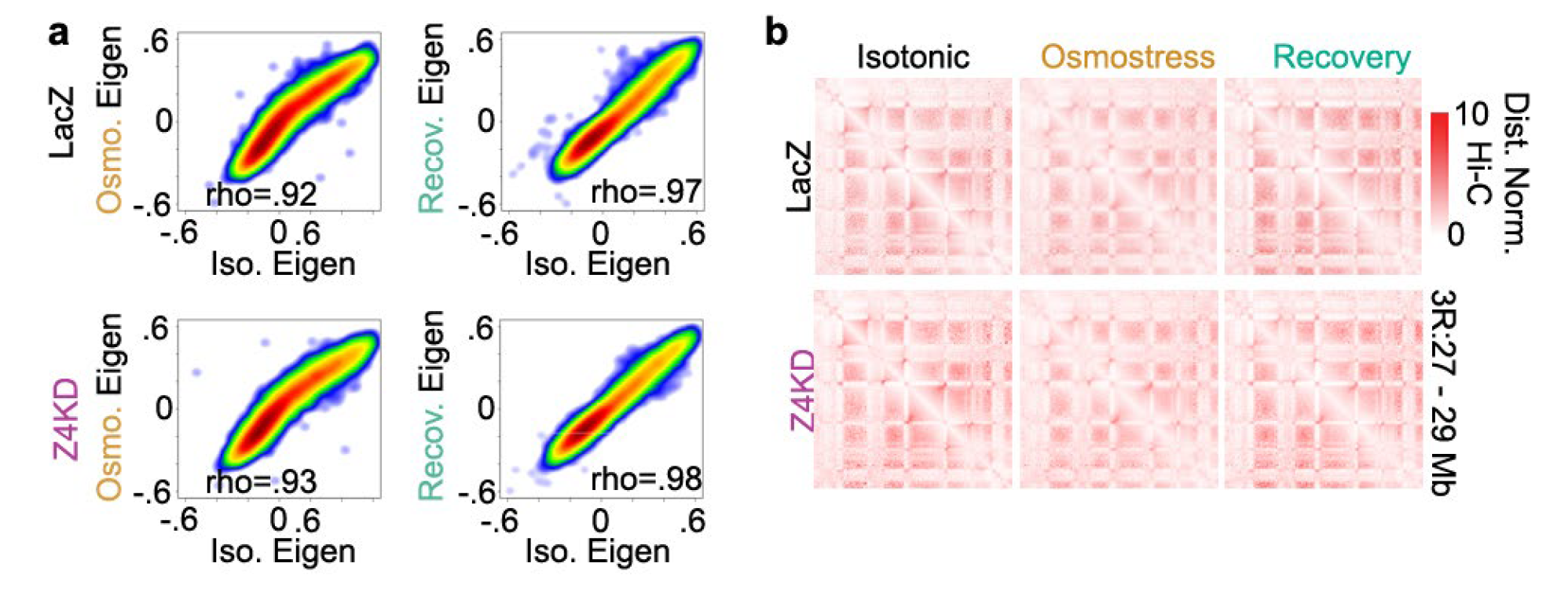
Extended Fig. 2. a) Comparison of the eigenvector between isotonic and either osmostress (left) or recovery (right) conditions in dsLacZ and Z4KD cells. Spearman rho is shown for each comparison. b) Example of the A/B compartment checkerboard pattern found under isotonic, osmostress, and recovery conditions in dsLacZ and Z4KD cells after distance normalization.

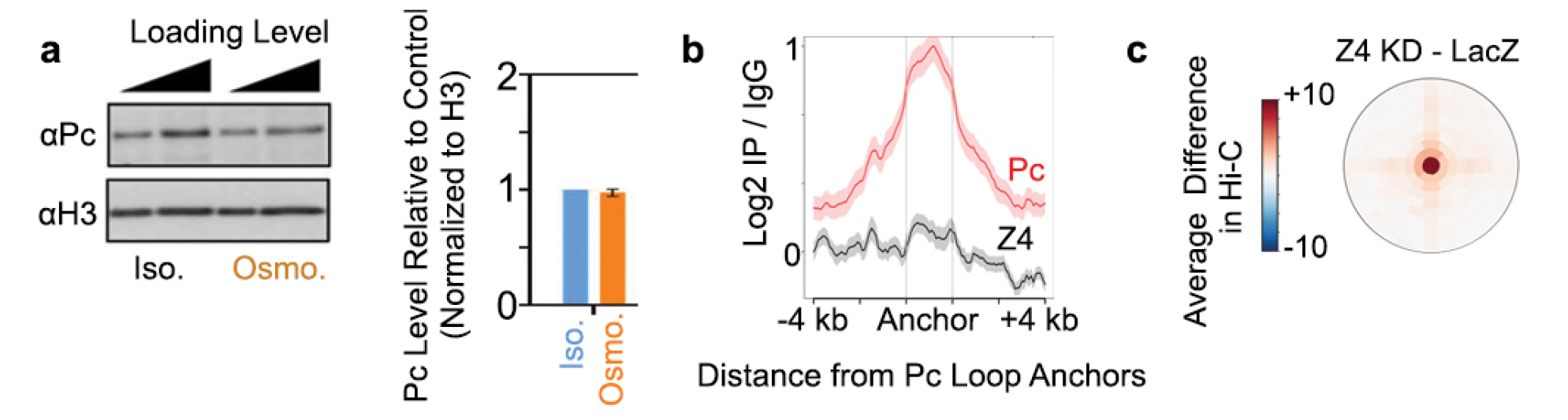
Extended Fig. 3. a) Left: WB for Pc on crosslinked chromatin in isotonic and osmostress conditions. H3 is shown as loading control. Right: Quantification of the results shown in the left. Results are the average of 3 independent experiments. Error bars are S.D. p-value > 0.05; two-tailed Student’s t-test. b) Average ChIP-seq signal relative to IgG for Pc and Z4 at Pc loop anchors. c) Average difference in Hi-C signal at Pc loops in Z4KD cells compared to dsLacZ cells. Distance normalized.

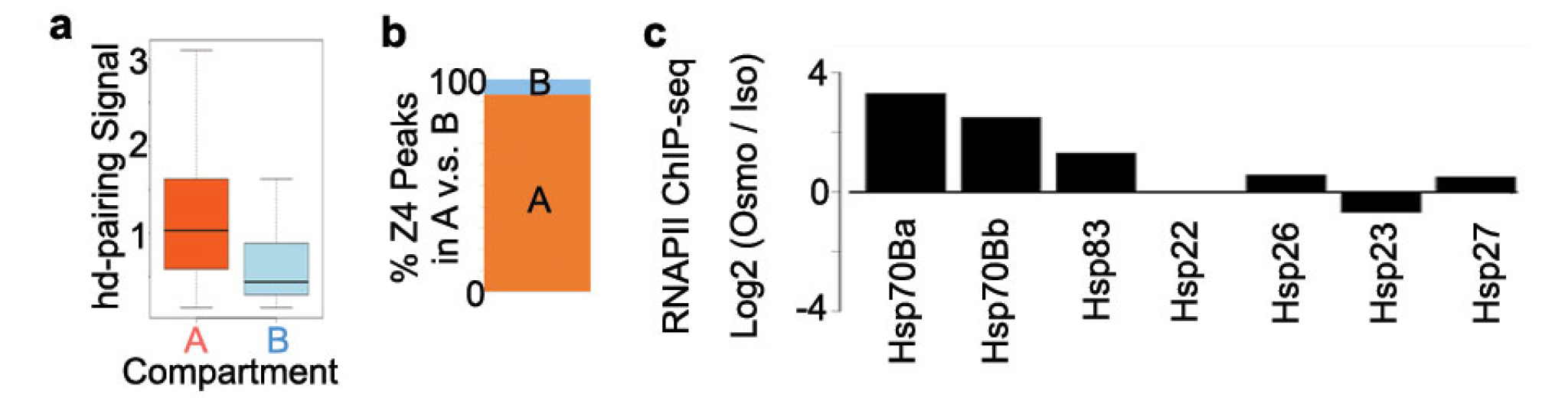
Extended Fig. 4. a) Boxplot of the hd-pairing signal in A and B compartment intervals. b) Percentage of Z4/CAP-H2 peaks found in A and B compartment intervals. c) Log2 change in RNAPII ChIP-seq signal for heatshock genes in osmostress compared to isotonic conditions.

**Extended Table 1.**
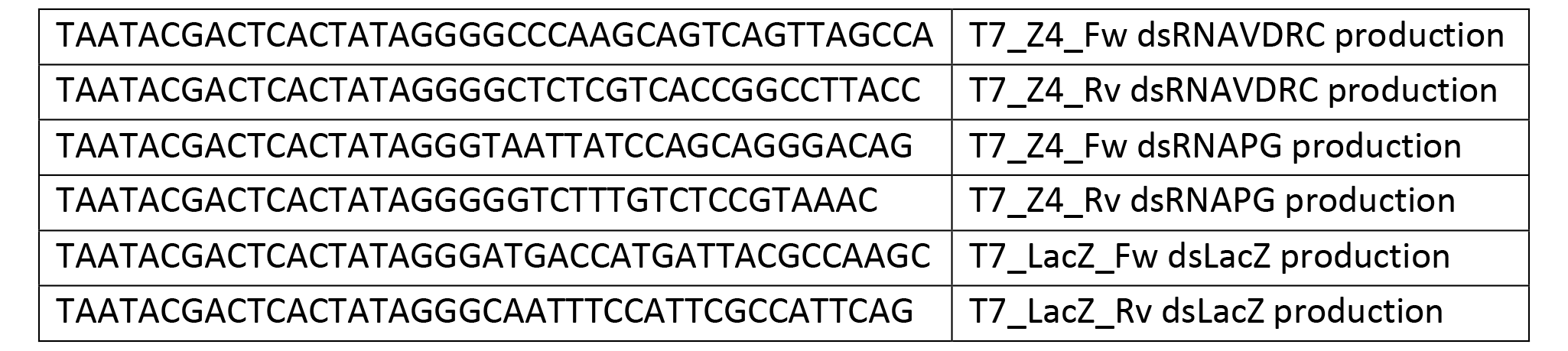
Primers used in RNAi experiments

**Extended Table 2.** Hi-C sequencing statistics

|  | **LacZ Isotonic** |  |  | **Z4 KD Isotonic** |  |  |
| --- | --- | --- | --- | --- | --- | --- |
|  | Rep1 | Rep2 | Combined | Rep1 | Rep2 | Combined |
| Total Pairs Processed | 345,840,775 | 347,659,645 | 693,500,420 | 374,987,680 | 355,965,025 | 730,952,705 |
| Unque Aligned Pairs | 153,070,165 | 151,208,184 | 304,278,349 | 162,910,007 | 160,579,466 | 323,489,473 |
| Trans Interactions | 4,526,989 | 3,984,105 | 8,511,094 | 3,756,076 | 2,044,990 | 5,801,066 |
| Cis Interactions | 96,653,019 | 91,701,048 | 188,354,067 | 94,052,304 | 94,906,705 | 188,959,009 |
| Cis short range | 38,475,765 | 36,865,869 | 75,341,634 | 39,293,537 | 45,750,172 | 85,043,709 |
| Cis long range | 58,177,254 | 54,835,179 | 113,012,433 | 54,758,767 | 49,156,533 | 103,915,300 |
| Deduplicated Valid Pairs | 101,180,008 | 95,685,153 | **<u>196,865,161</u>** | 97,808,380 | 96,951,695 | **<u>194,760,075</u>** |
|  | **LacZ Osmostress** |  |  | **Z4 KD Osmostress** |  |  |
|  | Rep1 | Rep2 | Combined | Rep1 | Rep2 | Combined |
| Total Pairs Processed | 362,792,625 | 351,116,114 | 713,908,739 | 395,902,828 | 298,636,038 | 694,538,866 |
| Unque Aligned Pairs | 156,590,039 | 147,171,365 | 303,761,404 | 169,195,596 | 128,540,359 | 297,735,955 |
| Trans Interactions | 11,086,178 | 8,101,687 | 19,187,865 | 8,791,393 | 3,754,509 | 12,545,902 |
| Cis Interactions | 98,154,685 | 93,763,281 | 191,917,966 | 109,401,762 | 72,125,876 | 181,527,638 |
| Cis short range | 20,230,249 | 20,107,535 | 40,337,784 | 24,901,764 | 25,906,459 | 50,808,223 |
| Cis long range | 77,924,436 | 73,655,746 | 151,580,182 | 84,499,998 | 46,219,417 | 130,719,415 |
| Deduplicated Valid Pairs | 109,240,863 | 101,864,968 | **<u>211,105,831</u>** | 118,193,155 | 75,880,385 | **<u>194,073,540</u>** |
|  | **LacZ Recovery** |  |  | **Z4 KD Recovery** |  |  |
|  | Rep1 | Rep2 | Combined | Rep1 | Rep2 | Combined |
| Total Pairs Processed | 295,446,488 | 329,486,272 | 624,932,760 | 363,329,856 | 368,351,970 | 731,681,826 |
| Unque Aligned Pairs | 131,822,342 | 142,808,082 | 274,630,424 | 156,265,124 | 172,396,010 | 328,661,134 |
| Trans Interactions | 3,822,566 | 3,566,246 | 7,388,812 | 4,666,838 | 2,653,870 | 7,320,708 |
| Cis Interactions | 85,752,466 | 85,457,721 | 171,210,187 | 98,497,050 | 105,222,510 | 203,719,560 |
| Cis short range | 29,147,540 | 30,009,046 | 59,156,586 | 32,837,223 | 47,547,357 | 80,384,580 |
| Cis long range | 56,604,926 | 55,448,675 | 112,053,601 | 65,659,827 | 57,675,153 | 123,334,980 |
| Deduplicated Valid Pairs | 89,575,032 | 89,023,967 | **<u>178,598,999</u>** | 103,163,888 | 107,876,380 | **<u>211,040,268</u>** |

